# Cell growth kinetics and accumulation of secondary metabolite of *Bletilla striata* Rchb.f. using cell suspension culture

**DOI:** 10.1101/698258

**Authors:** Yinchi Pan, Delin Xu, Shiji Xiao, Zhongjie Chen, Surendra Sarsaiya, Shebo Zhang, Yanni ShangGuan, Houbo Liu, Lin Li, Jishuang Chen

**Author notes:** Corresponding Author: Lin Li,; Jishuang Chen. Equal contribution (Y. Pan), (D. Xu).

## Abstract

*Bletilla striata* (Orchidaceae) is a well-recognized endangered medicinal plant due to inadequate natural reproduction with high market worth. To evaluate the cell growth kinetics and accumulation of secondary metabolites (SMs), the cell suspension culture is proved to be a valuable approach for acquiring the high yield of medicinal parts. An effective cell suspension culture for obtaining *B. striata* cell growth and its SMs was *in vitro* induction of callus from *B. striata* seeds. The cell growth kinetics and accumulation of SMs were analyzed using the mathematical model. Results cell growth kinetic model revealed that the growth curve of *B. striata* suspension cells was curved as sigmoid shape, indicating the changes of the growth curve of suspension cells. Improved Murashige and Skoog cell growth medium was the utmost favorable medium for *B. striata* callus formation with the highest cell growth during the stationary phase of cultivation period, the cell growth acceleration was started after 7 days and thereafter gradually decrease at 24 day and then reached to highest at 36 day of cultivation period in both dry weight and fresh weight. The coelonin concentration was peak during exponential growth stage and decreased afterward at the stationary phase in the cell suspension culture. The maximum content of coelonin (about 0.3323 mg/g cell dry weight) was observed on the 18th day of the cultivation cycle while the dactylorhin A and militarine reached highest at 24 day, and *p*-hydroxybenzyl alcohol at 39 day. This investigation also laid a foundation for multi-mathematical model to better describe the accumulation variation of SMs. The production of SMs had shown great specificity during cells growth and development. This research provided a well-organized way to more accumulation and production of SMs, on scale-up biosynthesis in *B. striata* cell suspension culture.

## 1. Introduction

*Bletilla striata* is a perennial herb of Orchidaceae, which is an important traditional Chinese herbal medicine recorded in the pharmacopoeia of all previous dynasties. It is sweet, and slightly cold with prominent effects of healing muscles and stopping bleeding. Hence is ideal for treating traumatic bleeding^[1]^. At present, due to the limitation of the traditional artificial breeding methods, the production of *B. striata* tuber and its medicinal active ingredients is difficult to meet the market demand^[2]^. *B. striata* has turn out to be an endangered class with the falling down wild plant properties due to over utilization in current periods. For the sustainable progress and comprehensive consumption of *B. striata*, it is essential to recognize its cell growth dynamics with the accumulation of SMs. However, until now, limited approaches have been developed on these perspectives. In addition, the quality and yield of medicinal substances have been limited because of the deprivation of cultivated variations in the course of long-period cultivation. Hence, it is needed to improve the medicinal properties with upright quality, more yield, and important secondary metabolites, for sustainable convention of medicinal resources.

With this view, the plant cell suspension culture technology can achieve artificial control to provide the optimal conditions for cell growth, differentiation and accumulation of SMs, therefore, we can efficiently promote cell proliferation and directionally induce the synthesis and accumulation of secondary metabolites^[3]^. This technology has become the most promising biosynthetic method for producing secondary metabolites from plant cells. Cell suspension culture systems of various medicinal plants have been established at national and global level, but studies on cell suspension culture and detection of secondary metabolites of *B. striata* have rarely been reported^[4,5]^.In the early stage, our research group has isolated, purified and identified a variety of secondary metabolites from the tuber of *B. striata*^[6,7]^. However, it is still unknown whether these secondary metabolites are also present in *B. striata* suspension culture cells.

On the basis of baseline research of our group, *B. striata* seeds have successfully been used to induce callus and establish a rapid propagation system^[8]^. Based on the induction and proliferation of *B. striata* callus, in this paper, we used an induced loose callus as the initial material to establish an optimized cell suspension culture system, and drew the growth curve. The changes of the accumulation of four major SMs, p-hydroxybenzyl alcohol, dactylorhin A, militarine and coelonin, were also detected. These laid a foundation for the further development of *B. striata* cell suspension culture bioreactors, as well as genetic improvement, regulation of cell proliferation, SMs production, and improvement of the efficiency of producing pharmaceutical important ingredients by using *B. striata*.

## 2. Materials and methods

### 2.1 Experimental materials

The loose callus induced by mature seeds of *B. striata* was used and collected from the Zhengan, Guizhou, China. The callus was then inoculated on Murashige and Skoog (MS) medium supplemented with 6-Benzylaminopurine (6-BA) 1.0 mg/L+2,4-Dichlorophenoxyacetic acid (2,4-D) 3.0 mg/L+30 g/L sucrose+7 g/L agar powder and subcultured in the dark at 25°C. After 2 generations of about 30 days, callus with good growth, loose texture and uniform growth was selected as the explant of liquid suspension culture.

HPLC grade methanol for Burdick & Jackson ACS/HPLC was procured from Honeywell, USA, and the standards dactylorhin A (CAS: 256459-34-4), militarine (CAS: 58139-23-4), coelonin (CAS: 82344-82-9) from ChemFaces Corp. The standard p-hydroxybenzyl alcohol (CAS: 623-05-2) was purchased from Chengdu Ruifensi Biotechnology Corp., Ltd., China. The quality score of the reference substance was more than 98%.

### 2.2 Instruments

BL-100A autoclave and GZX-9146MBE electric drying oven were purchased from Shanghai Boxun Medical Bio Instrument Co., Ltd., China; CJHS ultra-clean workbench from Tianjin Taisite Instrument Co., Ltd., China; Agilent 1260 HPLC, DAD UV detector, ChemStation chromatography workstation from Agilent Company, United States; BT125D analytical balance 1/100,000 from Sartorius Company, Germany.

### 2.3 Construction of cell suspension culture system

The previously obtained loose, tender yellow callus of *B. striata* was inoculated into a 35 mL liquid medium (MS+6-BA 1.0 mg/L+2,4-D 3.0 mg/L+30 g/L sucrose) in a 100 mL flask with pH 5.8 and 1.0g per bottle (fresh weight). After inoculation, it was placed in a rotary shaker, with shaking speed at 120 r•min^-1^ at 25°C temperature under dark culture condition.

### 2.4 Determination of dry weight, fresh weight and growth curve

Based on the culture conditions as stated above, the fresh and dry weight were measured at every 3 days intervals after inoculation upto the 45 days taken for one cycle. During the study, the flask containing the suspension cell fluid on the shaker was removed, then shaken and filtered until no droplets formed. The fresh weight was obtained and then dried in an oven at 50°C to a constant weight. The dry weight was also measured. From each sample points, three samples were collected and measured at three times. The fresh and dry weights were recorded respectively, and the growth curve of the cells was plotted with the culture time as the abscissa and the fresh and dry weights as the ordinate.

### 2.5 Cell growth curve modeling method

According to the data of fresh and dry weights of suspension cells, the growth curve was plotted and analyzed by Logistic, Boltzmann and DoseResp with Origin 9.1 software. The best fitted model was determined by the Fitting value (R^2^), and the F value analyzed by ANOVA^[9]^. The selected function model was used to detect the acceleration rate of cell growth to analyze the proliferation of cells.

### 2.6 Extraction and detection of secondary metabolites from cells

#### 2.6.1 Preparation of test solution and chromatographic conditions

According to the sampling method of 2.4, the callus of different growth stages was used to detect the content of four target secondary metabolites. The sampling was repeated 3 times at every 3 days intervals for each detection point. The callus was filtered, dried, crushed, and passed through the sieve (Φ200×50mm). Approximately 0.20 g of *B. striata* cells were accurately weighed, mixed with 100 ml of 70% methanol water for 2 hours into the condensation reflux extraction, and centrifuged to obtain an extract liquid. The recover extract liquid was dried under reduced pressure, dissolved in an appropriate amount of 70% methanol water, and then transferred to a 5 mL volumetric flask for use as a test solution. HPLC detection conditions were as follows; Column: Dubhe C_18_ (250 mm × 4.6 mm, 5 μm); mobile stage: methanol (A), ultrapure water (B); flow rate: 0.8 mL / min; column temperature: 25 °C; detection wavelength: 225 nm^[10]^; injection volume: 20 μL. The changes of volume fraction during the gradient elution process is as follows: mobile stage methanol (A) - ultrapure water (B), gradient elution (0-10 min, 20%-25% A; 10-25 min, 25 %-50% A; 25-35 min, 50%-50% A; 35-50 min, 50%-100% A; 50-60 min, 100%-100% A).

#### 2.6.2 Study on the linear relationship of the standard curve

A suitable concentration of p-hydroxybenzyl alcohol, dactylorhin A, militarine, coelonin were prepared for standard solution, and diluted with methanol to prepare a series of mass concentrations (0.25, 0.2, 0.15, 0.1, 0.05, 0.025 mg/mL for p-hydroxybenzyl alcohol, 1.0, 0.8, 0.6, 0.4, 0.2, and 0.1 mg/mL for militarine, 0.1, 0.05, 0.025, 0.01, 0.005, and 0.0025 mg/mL for coelonin). The detection was done under the standard chromatographic conditions as described in the section **2.6.1**.

#### 2.6.3 Validation of HPLC method

##### 2.6.3.1 Precision experiment

The standard solutions as described in the section **2.6.2** were precisely taken, and 20 µL set for injection volume. The sample was detected under the chromatographic conditions of **2.6.1** and repeated for 6 times.

##### 2.6.3.2 Stability experiment

Sample solution (as described in **2.6.1)** was injected at 0, 3, 6, 9 and 12 hours after preparation. Peak areas of p-hydroxybenzyl alcohol, dactylorhin A, militarine and coelonin were recorded, and the injection volume was 20 µL.

##### 2.6.3.3 Repeatability of experiment

Take 5 *B. striata* cell clusters, each of which was weighed 0.20 g. The samples were prepared according to the method under **2.6.1**. All samples were injected under the same chromatographic conditions. The injection volume was 20 µL. The peak areas were recorded and calculated the the mass fractions of p-hydroxybenzyl alcohol, dactylorhin A, militarine and coelonin. The relative standard deviation (RSD) was calculated in according to the formula.

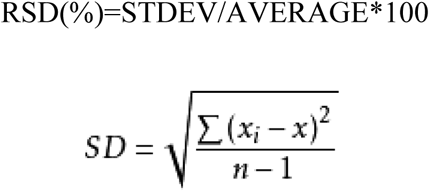

##### 2.6.3.4 Sample recovery analysis

Nine *B. striata* suspension cultured cell clusters with known secondary metabolites were accurately weighed, each of which was weighed 0.02 g, and precisely added to three reference solutions with low, medium and high mass concentrations (80%, 100%, 120% of the original sample, respectively). There were 3 reference substances in each mass concentration. The recovery rate and RSD of each components were calculated.

### 2.7 Modeling method for cumulative curve of secondary metabolites

The data of the cumulative number of secondary metabolites of *B. striata* suspension cells were fitted by a variety of functional models under the “Nonlinear Curve Fit” in Origin 9.1 software and determined the best fitted model by the fitness value (R^2^).

## 3 Results

### 3.1 Growth curve of suspension cultured cells

The growth curve of suspension culture cells was plotted with the fresh and dry weight as the indicators. As shown in Figure 1, it was found that the growth curves of the two different growth indicators was basically the same and both were “sigmoid” type.

**Figure 1.**
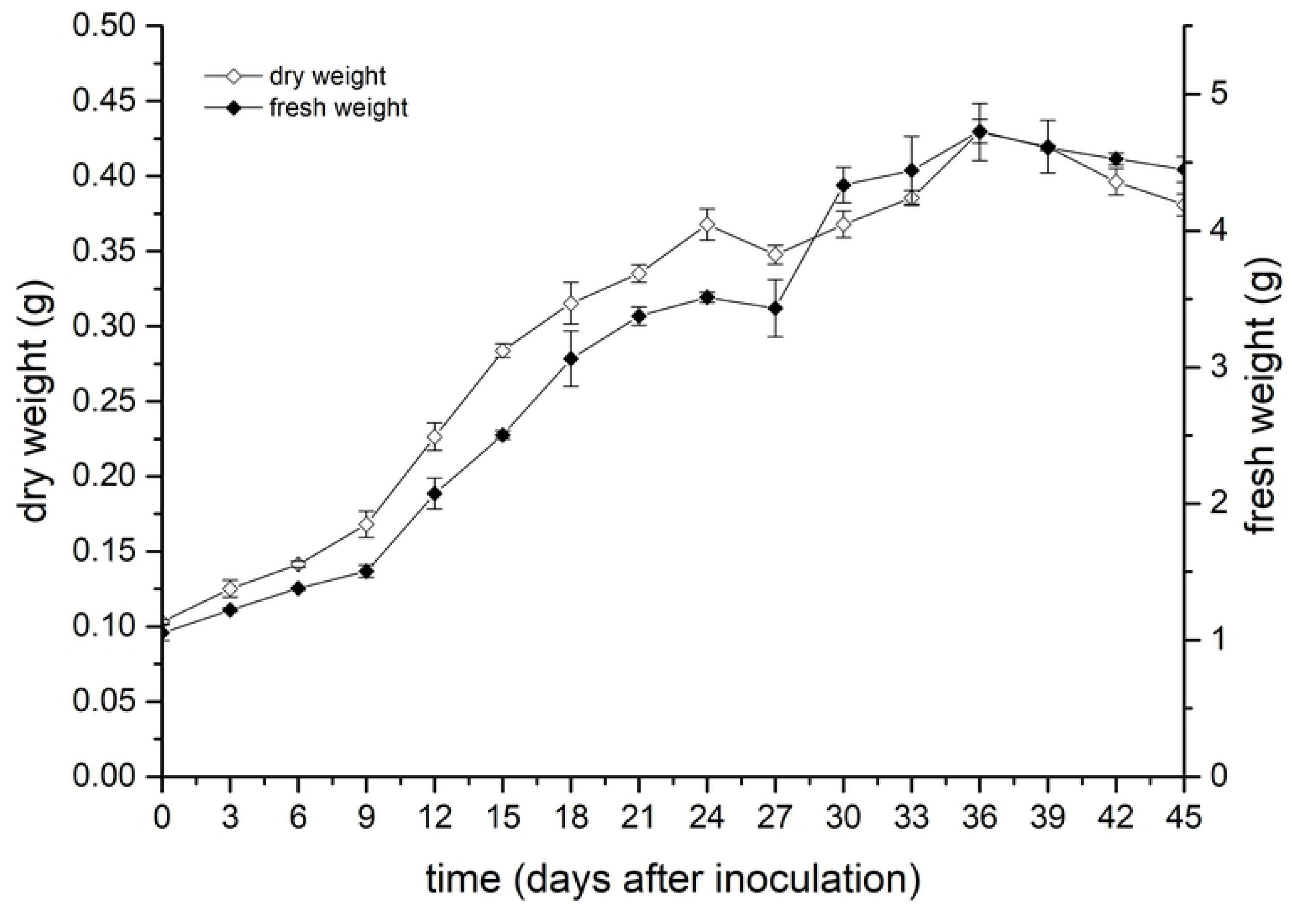
Growth curve of fresh and dry weights of suspension cells in culturing days.

### 3.2 Growth curve and kinetic characteristics

#### 3.2.1 Functional model of growth curve

The mathematical models were fitted and analyzed for the change of fresh and dry weight of suspension culture cells. The results are shown in Table 1.

**Table 1.**
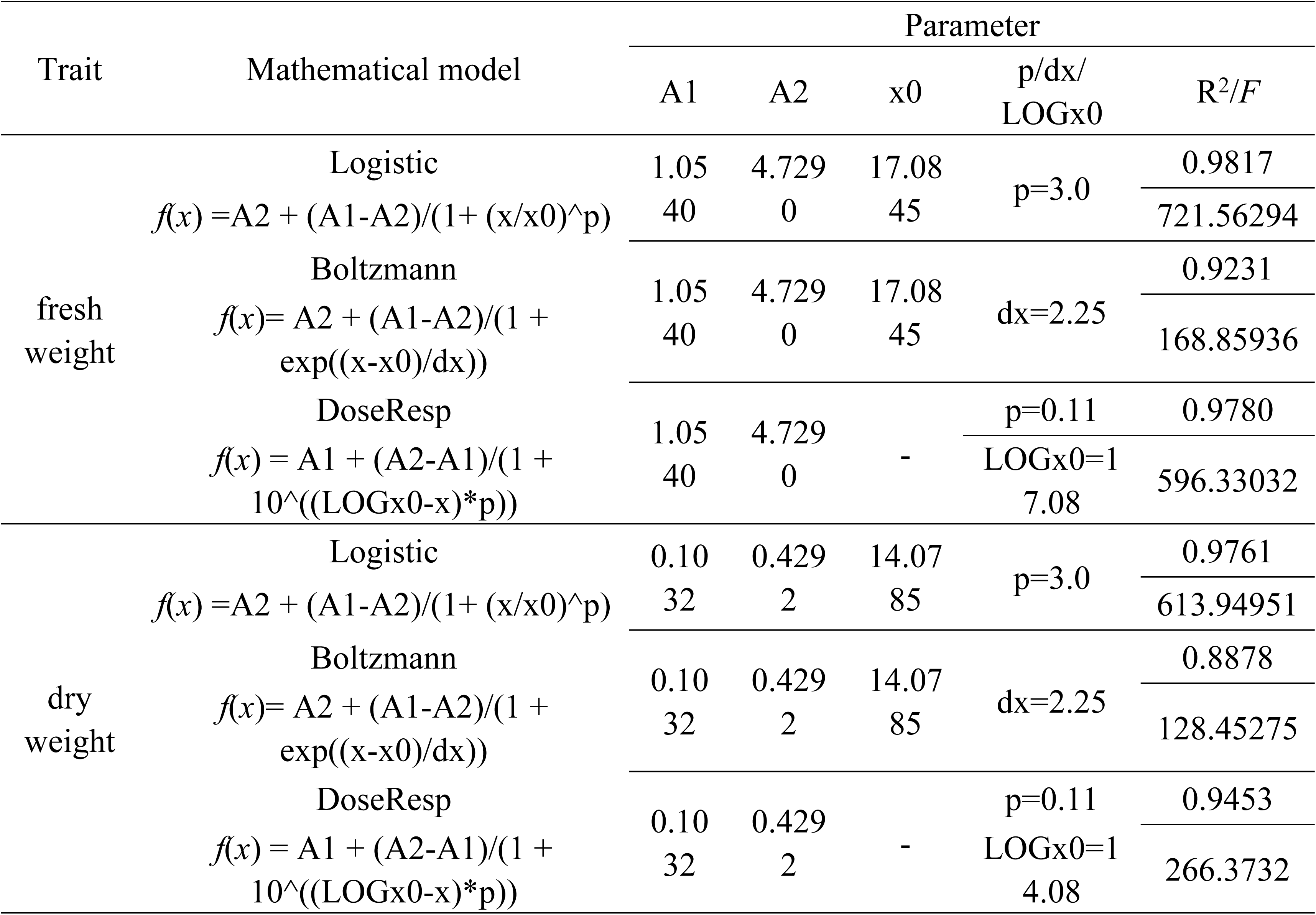
Mathematical modeling results of the growth curve of fresh weight and dry weight of suspension culture cells.

As shown in Table 1, the coefficient of determination of the logistic, Boltzmann and DoseResp fitting equations for the fresh weight and days were 0.9817, 0.9231, and 0.9780, respectively. The coefficient of determination of the three fitting equations for dry weight and days were 0.9761, 0.8878 and 0.9453, respectively. According to the fitting results, the Logistic model was more suitable to describe the growth of biomass in suspension cells of *B. striata*. After a further significance test, the F values of Logistic equation (721.56294 and 613.94951) were also higher than Boltzmann equation and Dose Resp equation, indicating that the curve of Logistic equation is more consistent with the experimental data. Logistic equation was further used to plot the curve of the fresh and dry weight growth of suspension cultured cells (Fig. 2).

**Figure 2.**
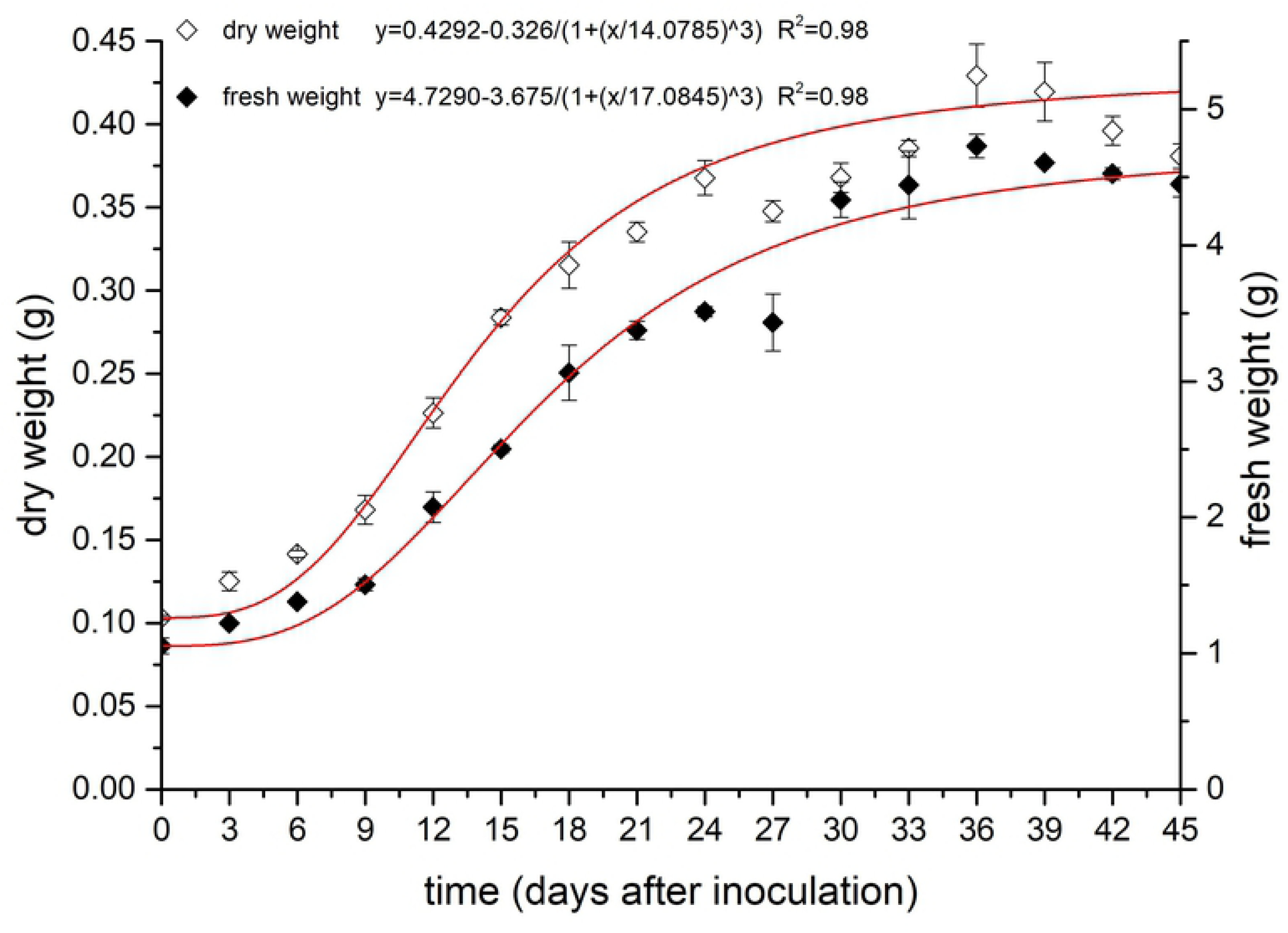
Growth kinetics curves of fresh and dry weights of suspension cells.

The results showed that the whole culture cycle could be divided into six stages: lag stage, exponential stage, linear stage, deceleration stage, stationary stage and recession stage. Among them, the lag stage was found on 0 to 6 day. During this period, the fresh and dry weight of suspension culture cells were changed slowly, indicating that the cells gradually adapted to the environment. During the exponential stage (between 6-12 day), the cell growth rate was gradually increased and reached to the maximum. The growth rate of callus was gradually stable during the linear stage of 12-24 day, and the change of fresh and dry weight was linearly correlated with time. The deceleration stage was lasted from 24 to 36 day, during which the cell growth rate gradually decreased. At 36 day, the fresh and dry weight of cells were reached the maximum. While they did not change significantly during the stationary stage of 36-39 day while these two weights were beginning to decrease after 39 day.

#### 3.2.2 Cell growth curve function model analysis

The Logistic function was presented the cumulative amount of cell growth. In order to better understand the changes in suspension culture cell growth, we performed first-order derivative (cell growth rate) and second-order derivative (cell growth acceleration) of the simulated Logistic function and plotted the results of the derivation, as shown in Figure 3. It is reflected from the figure that the growth rate of the cells was increased at initially. When it reached to its peak, it was affected to decrease gradually by various factors and the growth rate. The overall trend was first increased and then decreased. The extremums of the two curves showed that the growth acceleration of suspension cultured cells was reached the maximum on the 7 day and the cell growth rate reached the maximum on 13 to 14 day.

**Figure 3.**
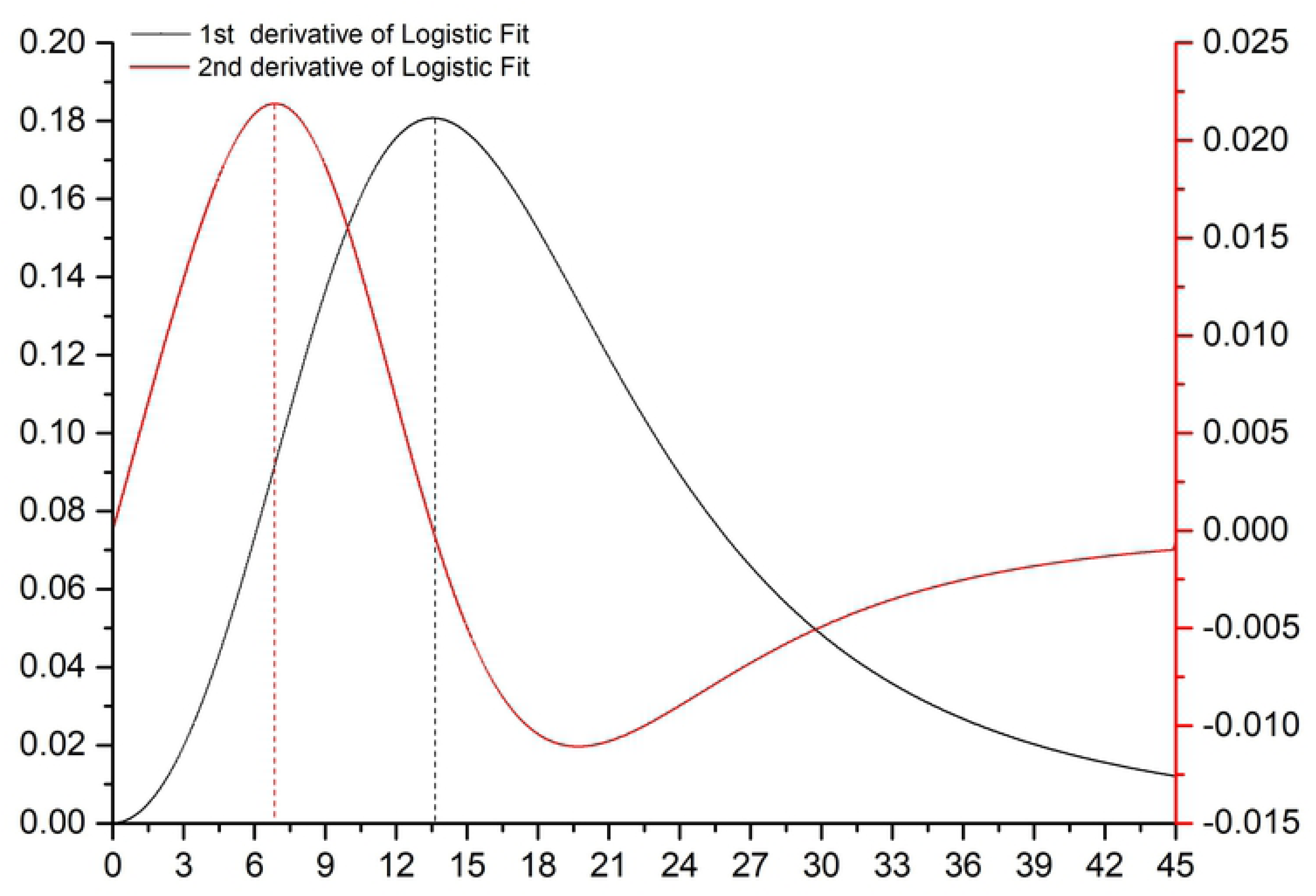
Function graphs of the first derivative and the second derivative of Logistic model.

### 3.3 Measurement of secondary metabolites accumulation in suspension cultured cells

#### 2.3.1 Investigation of specificity

The chromatograms samples were tested with reference substances, filtrate of suspension culture cells and blank solution are shown in Fig. 4. The results shown that the absorption peaks of the sample was tested at the same conditions as the reference substance, and the blank solution for no interference. Moreover, it showed that there was no obvious number of secondary metabolites left in the cell culture medium, so the samples were accurately reflected the accumulative amount of secondary metabolites in suspension culture cells.

**Figure 4.**
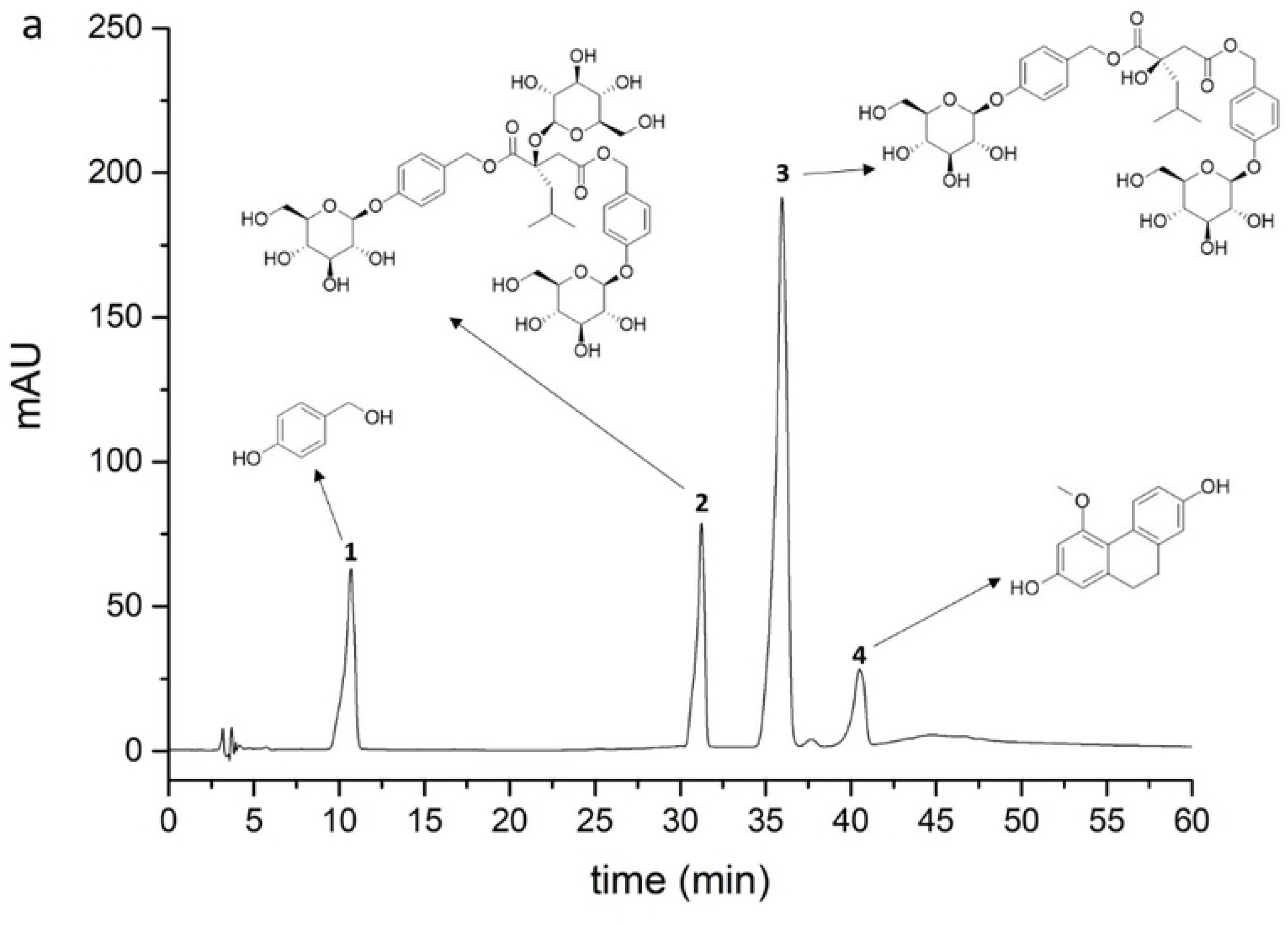
HPLC chromatograms of various constituents. a. Test samples. b. Reference substances. c. the filtrate of suspension cultured cells of *Bletilla striata.* d. Blank solution (70% methanol). 1. p-hydroxybenzyl alcohol. 2. dactylorhin A. 3. militarine. 4. coelonin.

#### 2.3.2 Investigation of linear relations

The standard curves were drawn with the mass concentration of p-hydroxybenzyl alcohol, dactylorhin A, militarine and coelonin as the abscissa (X) and the four secondary metabolites corresponding peak area as the ordinate (Y_1-4_). The regression equations were Y_1_= 91387X-169.99 (*R*^2^=0.9994), Y_2_= 36075X-712.50 (*R*^2^=0.9996), Y_3_= 24341X-224.88 (*R*^2^=0.9993) and Y_4_= 69896X-142.88 (*R*^2^=0.9994).

#### 2.3.3 Validation of the methodology of HPLC

The results of precision test showed that RSD of peak areas of p-hydroxybenzyl alcohol, dactylorhin A, militarine and coelonin were 0.83%, 0.40%, 1.36%, 1.24%, respectively while RSD of retention time were 0.09%, 0.08%, 0.07%, 0.16%, respectively. The results of stability test showed that the RSD of peak areas of the four components were 0.71%, 1.73%, 1.86% and 2.80% respectively, while RSD of retention time were 0.86%, 0.50%, 0.76% and 1.61% respectively, indicating the solution had good stability within 12 hours. The results of repeatability experiment showed that the RSD of the four components were 2.19%, 2.54%, 0.78% and 2.00%, respectively, which indicated good repeatability. The recovery experiment of four secondary metabolites results were 1.59%, 1.85%, 1.24% and 1.98% showed that the method had good accuracy.

### 3.4 Kinetic characteristics of secondary metabolites accumulation in suspension system

The accumulation of the four secondary metabolites in cells of different growth stages were calculated, and plotted as curves. The mathematical models were used to describe the changes of the accumulation of the four chemicals, as shown in Table 2. The results showed that the changes of secondary metabolites in suspension cultured cells were more complicated than that of cell growth. A single mathematical model could not well describe the measured data. Therefore, for the three secondary metabolites (dactylorhin A, militarine, coelonin), the multi-mathematical model and piecewise function were selected to describe the change of their cumulative amount, as shown in Figure 5.

**Table 2.**
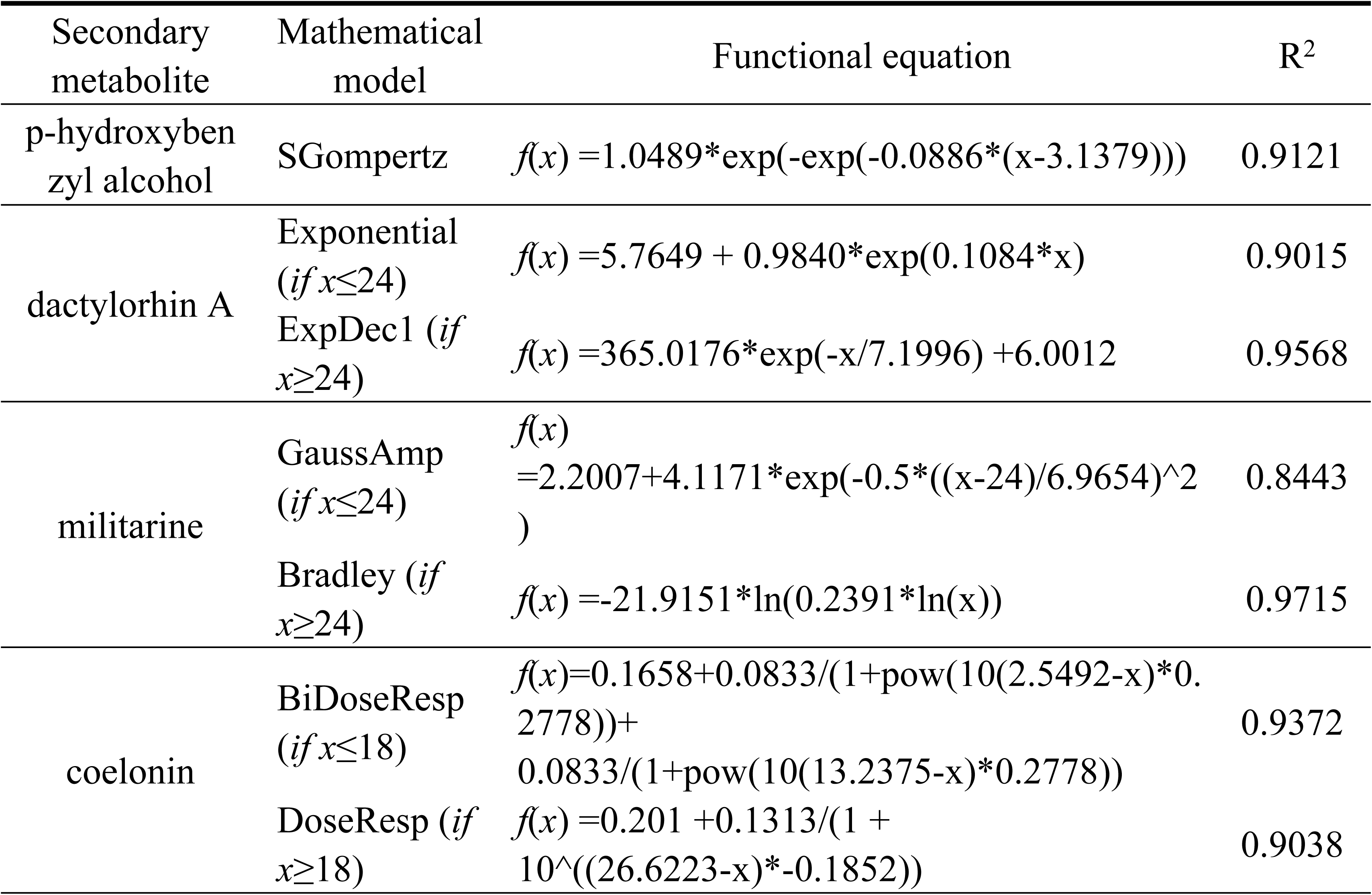
Mathematical model fitting results of four secondary metabolite content.

**Figure 5.**
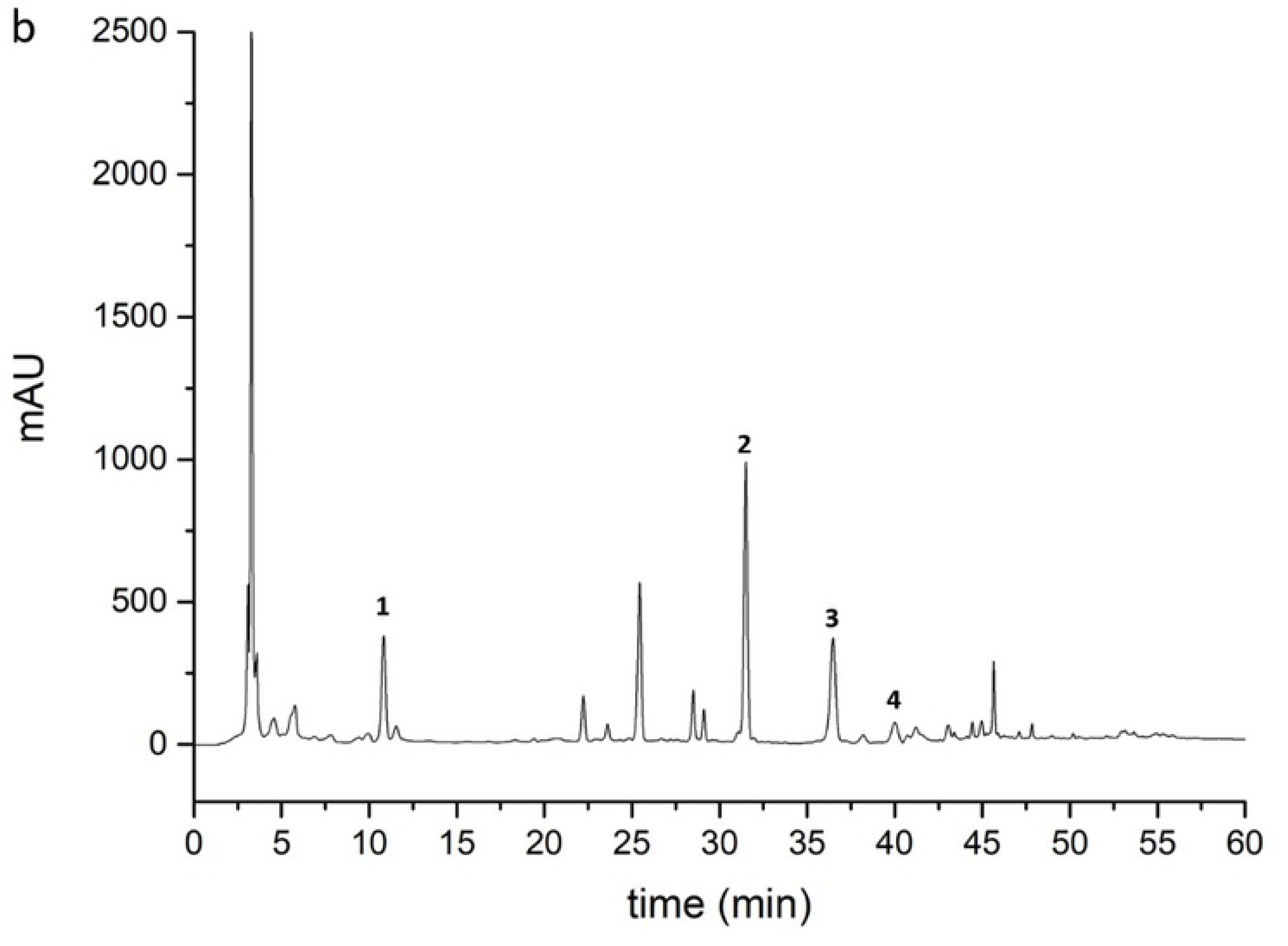
Content change and fitting curves of four secondary metabolites. a. p-hydroxybenzyl alcohol. b. dactylorhin A. c. militarine. d. coelonin.

According to the figure, the accumulation of p-hydroxybenzyl alcohol was similar to cell growth, showing a gradual upward trend. The curve went smoothly, and the cumulant increased slowly at 24 day, and the accumulation reached the maximum at 39 day. There was a distinct maximum value in the cumulant curves of dactylorhin A, militarine and coelonin. The cumulative amount of dactylorhin A and militarine both reached the maximum at 24 day, and the cumulative amount of coelonin reached the maximum at 18 day.

## 4 Discussion and conclusion

Compared with plant cultivation, cell culture has the advantages of short growth cycle, easy separation of secondary metabolites and easy control of influencing conditions. Plant cell culture makes it easier to obtain specific natural products from medicinal plants by specifically inducing the synthesis of specific secondary metabolites. Therefore, using mathematical models to analyze cell growth and secondary metabolite synthesis and accumulation is important for revealing the synthesis mechanism of natural products such as secondary metabolic components, improving the yield of secondary metabolites and enhancing the medicinal value of medicinal plants^[11]^. In recent years, with the deepening of relevant researches, the medicinal value of various secondary metabolites in *B. striata* has been confirmed by scientific research. For example, p-hydroxybenzyl alcohol can increase the expression of genes encoding antioxidant proteins after focal cerebral ischemia, which can avoid oxidative stress and further damage on brain neurons^[12]^. The dactylorhin A and militarine can significantly improve memory impairment in mice which is caused by chemicals such as scopolamine, cycloheximide and alcohol^[13]^ while 2,7-dihydroxy-4-methoxy-9,10-dihydrophenanthrene (coelonin) has certain antiviral activity as a kind of dihydrophenanthrene compound. It is more meaningful to use plant cell liquid culture technology to regulate the synthesis of various secondary metabolites^[14]^.

In this study, we simulated the growth of *B. striata* suspension culture cells with a variety of mathematical function models, and properly simplified the complex environment in which a group of cells were grown together with the mathematical model of growth kinetics. From the results of function fitting, all the mathematical models could accurately describe the growth of cells at different growth stages and the changes of secondary metabolites accumulation. The growth of suspension culture cells are divided into six stages: lag stage, exponential stage, linear stage, deceleration stage, stationary stage and recession stage. Between 13 and 14 day, the cell growth rate was reached its maximum. After 39 day, the cell growth did not show a significant decline trend in the functional model, but according to the actual data and observation of the state of the culture cells, we found that the culture cells showed obvious browning, indicating the growth entered the recession stage. These phenomena were well presented in the functional model.

However, the changes of accumulation of secondary metabolites in the suspension culture cells of *B. striata* were complicated and diversified. The change in the cumulative amount of p-hydroxybenzyl alcohol basically followed the change in the growth of *B. striata* suspension culture cells, while the content changes of dactylorhin A, militarine and coelonin were not consistent with the growth of cells. By comparing the growth curve with the change curve of the accumulation of secondary metabolites, when the cell growth reached the end of the linear growth stage (12-24 day), the accumulation of secondary metabolites generally declined. At this stage, due to the reduction of nutrients in liquid medium, the environmental inhibition began to be greater than the cell growth, and secondary metabolism, as a non-essential substance of plant life, would gradually decrease. Dactylorhin A and militarine, as a glycoside compound, are both secondary metabolites and energy storage substances that actively decompose to maintain life in adversity, which may result in a decrease in secondary metabolites in cells^[15]^. Fermentation kinetics studies on microorganisms have shown that p-hydroxybenzyl alcohol itself can act as an inducer to induce fungi to increase extracellular polysaccharide production. In this study, we found that the curve of p-hydroxybenzyl alcohol accumulation gradually became smooth after 24 day, while dactylorhin A and militarine reached maximum values at the 24^th^ day. The effects of p-hydroxybenzyl alcohol on the key enzymes of polysaccharide synthesis in plant cells and some possible factors of synthesizing promotion need to be further studied. It is well known that the production of secondary metabolites by plants are the result of their adaptation to the ecological environment during the long-term evolution. The synthesis of these secondary metabolites is closely related to the growth state of plants and environmental conditions. Different types of secondary metabolites have different biosynthetic pathways in the different stages of plant growth.

Further studies are needed to control the metabolic pathways, key enzymes and key regulatory genes; so that we can improve the efficiency of suspension culture of *B. striata* cells and achieve efficiently directive induction of secondary metabolites by means of genetic engineering and metabolic engineering^[16]^. It also lays a theoretical and practical foundation for the follow-up researches on the growth, mutant induction, and secondary metabolite synthesis and bioreactor construction of *B. striata*.

## Acknowledgments

This research was financially supported by the National Natural Science Foundation of China (31560079, 31560102), the Scientific Project of Guizhou Province (QKH-ZY[2013]3002, QKH-LH[2014]7549, [2017]5733-001), Talents Promotion Project of Zunyi Medical University(JC2018-2-5(1)), the PhD Science Foundation of Zunyi Medical University (F-809), and the Talent Growth Project of Guizhou Education Department (KY[2017]194).

## Supporting information

**Figure S1.**
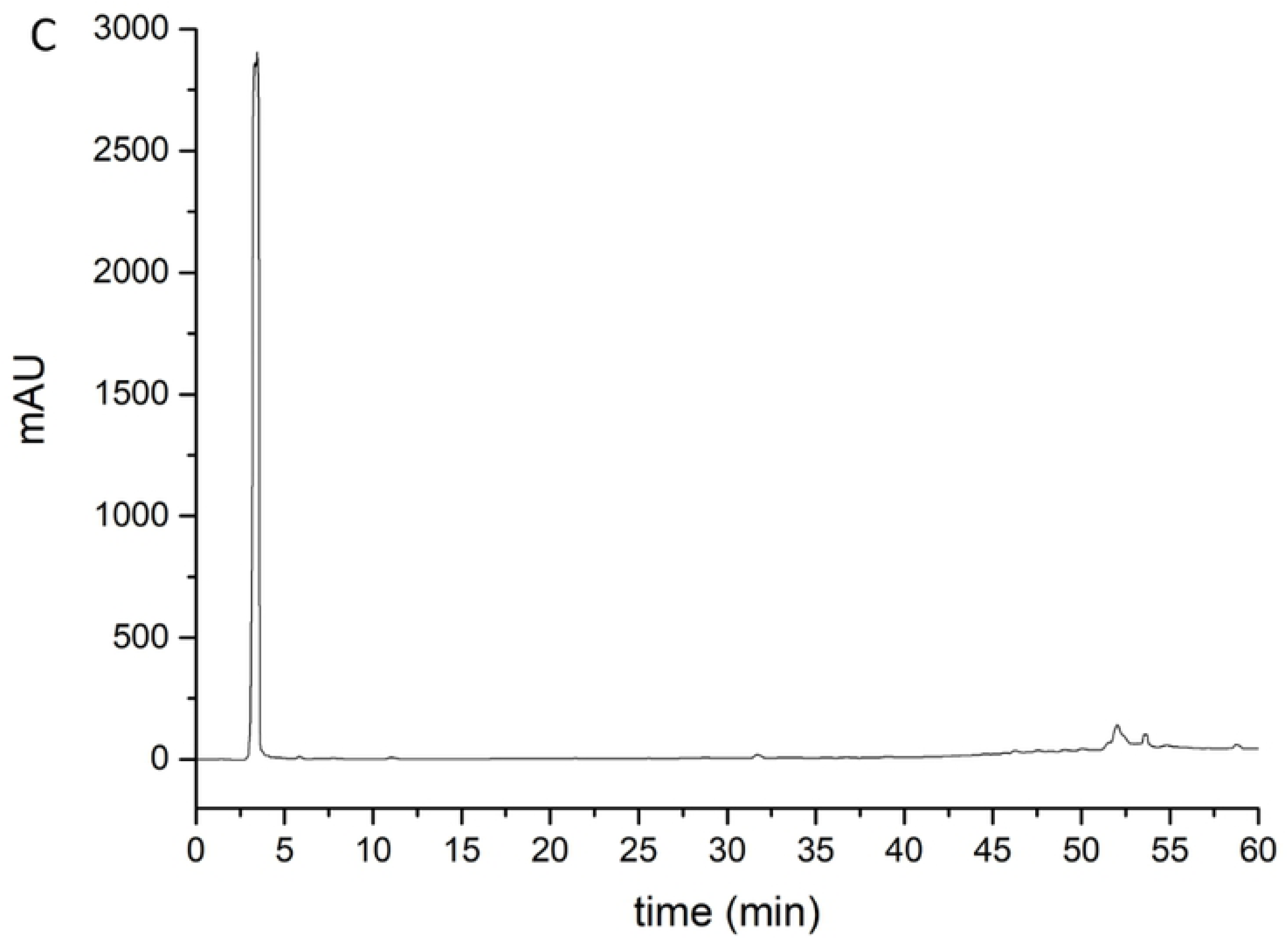
Morphology of callus in different growth periods. (.docx)

**Figure S2.**
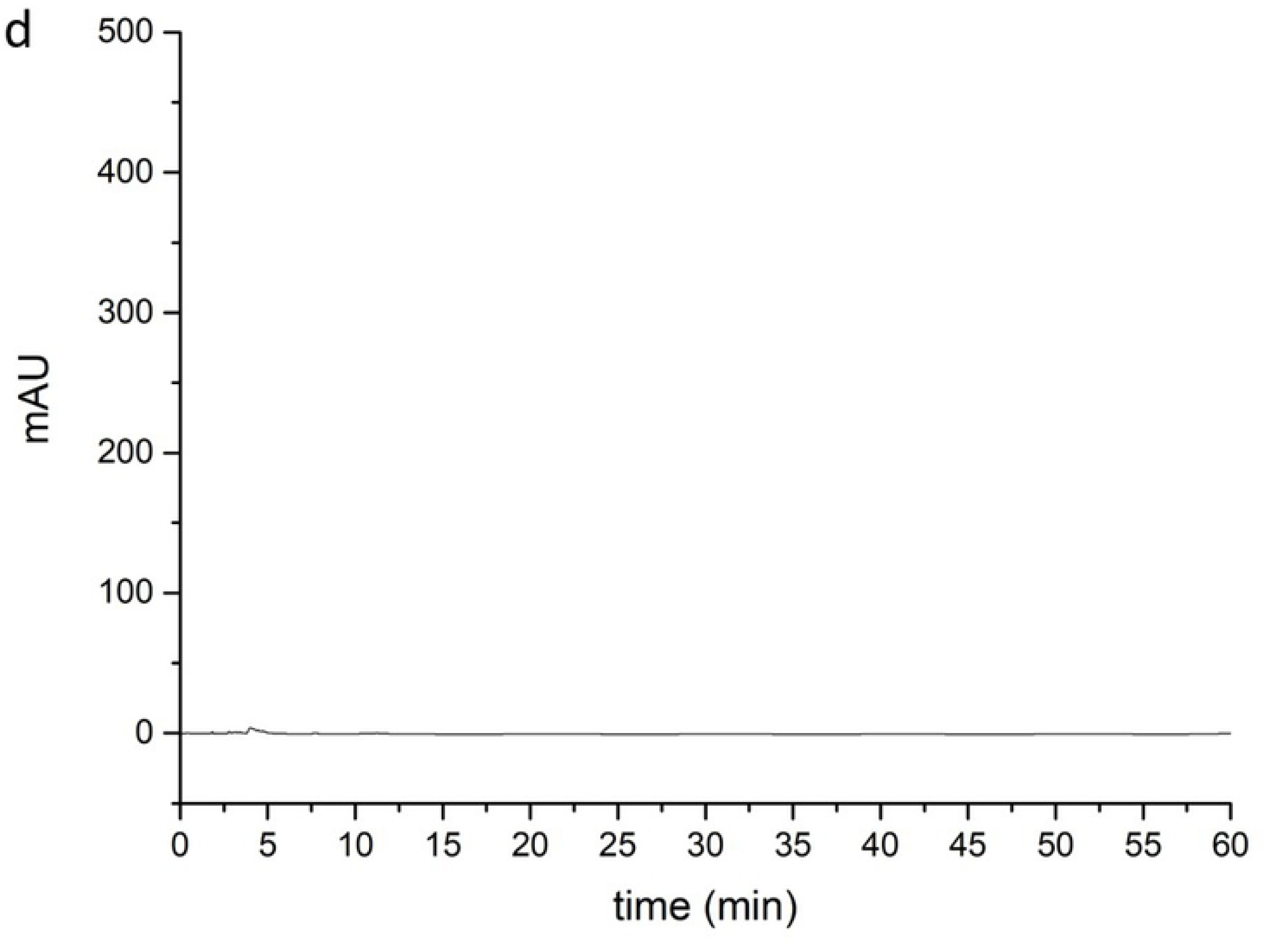
Growth states of suspension culture cell. (.docx)

**Figure S3.**
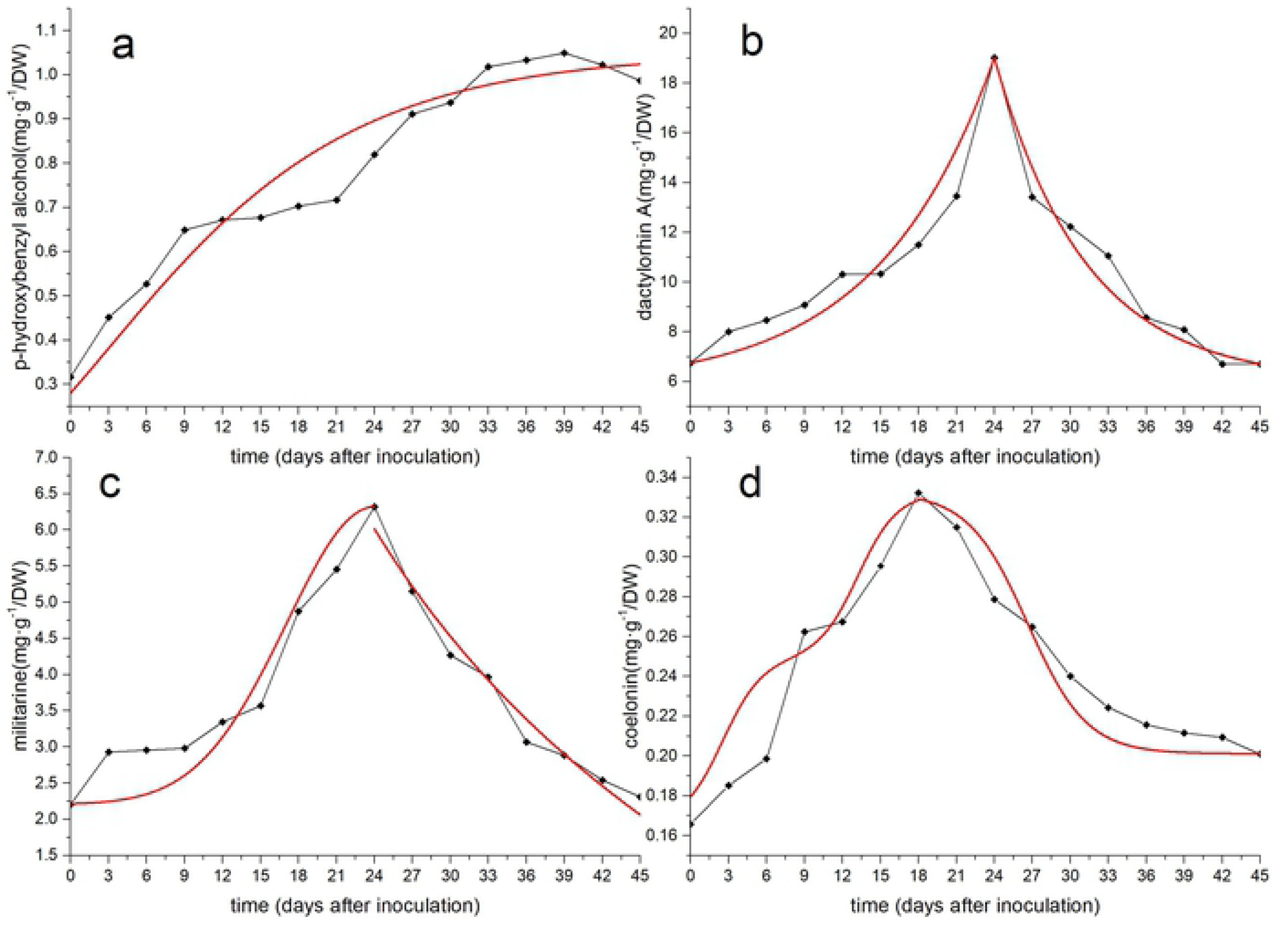
Standard curve of four secondary metabolites. (.docx)

Table S4. Examination results of precision. (.docx)

Table S5. Examination results of stability. (.docx)

Table S6. Examination results of reproducibility. (.docx)

Table S7. Results of recovery tests for tested chemicals. (.docx)

Table S8. Dry and fresh weight growth of suspension cultured cells in 45-day culture period. (.xlsx)

Table S9. Mean measurement of secondary metabolites. (.xlsx)

